# Relationship Between the P300 and Sequence Knowledge in a Changing Environment

**DOI:** 10.1101/2023.07.12.548771

**Authors:** Ko-Ping Chou, Tzu-Yu Hsu

## Abstract

The P300 amplitude has been linked to the processing of uncertain events. Studies have assumed that knowledge extracted from sequences of events corresponds to event probability. The relationship between the P300 and event uncertainty has been studied using the model-based analysis, in which the subjective expectancy of event probability is modeled and examined by using a quantity called “surprise.” However, other types of sequence knowledge exist, such as event transition probabilities, which comprises both event repetitions and event alterations. Whether the state of the environment affects the encoded sequence knowledge is not fully understood, and the type of sequence knowledge, event probability or event transition probability, that is encoded in the brain in a changing environment remains unknown. We determined whether fluctuations in the P300 are better explained by surprise based on a model of event probability or by surprise based on a model of event transition probability. Participants completed a two-choice response task in which a binary sequence was generated from a hidden Markov model. Reaction times indicated that behavior changed depending on the event transitions. The model-based analysis revealed that trial-by-trial P300 was better explained by surprise based on a model of event transition probability. Our results suggest that humans use the sequence knowledge of event transitions in a changing environment.

## Introduction

The P300 has been associated with the processing of unexpected events, and its magnitude is proportional to the uncertainty of the stimulus (<u>Sutton et al., 1965</u>). However, event-related potentials (ERPs) studies have increasingly suggested that the amplitude of the P300 not only reflects the stimulus probability but also individuals’ subjective experiences and expectancy of the underlying sequence structure (<u>Tueting et al., 1970</u>; <u>Squires et al., 1976</u>; <u>Squires et al., 1977</u>; <u>Duncan-Johnson and Donchin, 1977</u>; <u>Horst et al., 1980</u>; <u>Johnson and Donchin, 1980</u>; <u>Dochin et al., 1986</u>). This has led to the model-based analysis, in which computational models represent beliefs regarding events or rules that affect the decision-making process (<u>Corrado and Doya, 2007</u>). To determine whether computational models can represent beliefs, a concept in information theory called “surprise” can be quantified (<u>Strange et al., 2005</u>). The method of quantifying surprise is based on a model for quantifying event improbability that determines the amount of information conveyed by an event (<u>Strange et al., 2005</u>). The model-based analysis and surprise can be used to formally quantify the subjective probability of an event.

In addition to subjective beliefs regarding event probability (<u>Mars et al., 2008</u>; <u>Kolossa et al., 2013</u>), regularities in sequences strongly affect the magnitude of the P300 (<u>Dehaene et al., 2015</u>). Sequence knowledge enables the prediction of events and subsequent preparation. Two types of sequence knowledge are of particular interest: event probability and event transition probability. Event probability involves event frequencies, whereas event transition probability involves both event repetition and event alteration frequencies (<u>Meyniel et al., 2016</u>). Surprise in models of event probability is called “predictive surprise,” whereas that in models of event transition probability is called “predictive transition surprise.” Predictive surprise is more representative of central-parietal P3b, whereas predictive transition surprise is more representative of frontocentral P3a. Both types of sequence knowledge are encoded in the brain and related to the P300 (<u>Higashi et al., 2017)</u>.

Although studies have considered both event probability and event transition probability and demonstrated that the brain is sensitive to each type of sequence knowledge, most studies have adopted a block design (<u>Mars et al., 2008</u>; <u>Kolossa et al., 2013</u>; <u>Higashi et al., 2017</u>), whereby participants gradually adapt to a single sequence over time in blocks. The prediction of events may initially rely on event transitions, but given the stability of a sequence, event probability within blocks may also be useful because of lower computational demand. This phenomenon could be observed in Higashi et al. (2017) study, their results showed that model performance changed across different time courses. The model fitting accuracy of P300 amplitudes based on the event transition model was consistent across trials. More importantly, the performance of the event probability model was superior in later stages of a block. Thus, the block design ensures a stable-state environment, resulting in prediction being ceased at a certain time point. However, whether the state of the environment changes over time remains unknown, as does the type of sequence knowledge encoded in the brain when the changes occur.

Knowledge of an environment’s previous state becomes irrelevant as an environment changes. This knowledge must be discarded, and new knowledge based on the environment’s current state must be acquired, which shapes individuals’ beliefs regarding the environment (<u>Nassar et al., 2010</u>; <u>O’Reilly et al., 2013</u>). Studies on decision-making have demonstrated that decisions are not only guided by outcomes but are also modulated by higher-order statistics related to the environment, such as those indicating whether an environment is stable or changing (<u>Behrens et al., 2007</u>; <u>Meder et al., 2017</u>). Tracking an environment is crucial to decision-making (<u>Klein-Flügge et al., 2022</u>). The prediction of events also relies on determining the state of an environment. We investigated prediction-making in a changing environment and the type of sequence knowledge that the brain encodes. We hypothesized that the brain would encode the sequence knowledge of event transition probability, which creates a more detailed representation of an environment than does event probability. In addition, we hypothesized that the P300 would be better explained by surprise based on a model of event transition probability.

This study examined fluctuations in the P300 in a changing environment. Participants completed a two-choice reaction time (TCRT) task in which a binary sequence was generated from a hidden Markov model (HMM). HMM consists of two stochastic processes. The first identifies hidden state transitions by using a state transition matrix. The second controls the emission of observations in each state by determining emission probabilities. HMM enables a changing and probabilistic sequence to be created. To identify the type of sequence knowledge, we used the Bayes-optimal learning model from <u>Meyniel et al. (2016)</u> for two purposes. One purpose was to determine predictive transition surprise on the basis that participants learned the sequence knowledge of event transition probabilities, and the other was to determine predicted surprise on the basis of event probabilities. The model-based analysis was used to determine whether reaction times (RTs) and the P300 amplitudes reflect predictive surprise, predictive transition surprise, or both in a changing environment.

## Methods

### Participants

Thirty-eight participants aged between 21 and 37 years (20 women, average age: 26 years) participated in the experiment at Shuang-Ho Hospital, New Taipei City, Taiwan. Participants were recruited from local communities and had no history of neurologic or psychiatric disorders. They were not taking any psychotropic medications. All participants provided informed consent. This study was approved by the Taipei Medical University Joint Institutional Review Board (N201903148). All participants received 1,000 NTD as compensation for their time.

### Apparatus and Procedure

The experimental stimuli were presented on a display (58 cm × 53 cm, ViewSonic) with a 60-Hz refresh rate. The experiment was made using Psychopy (v3.1.5). Participants were seated 60 cm in front of the display, with their heads resting on a chin rest. The experiment was divided into two sessions, each consisting of a continuous sequence with 400 trials. The sequences differed between sessions. Each session was approximately 20 minutes, with a 5– 10-minute break between sessions.

The experiment was based on TCRT tasks in other studies (<u>Higashi et al., 2017</u>; <u>Meyniel et al., 2015</u>). A binary sequence was presented without feedback regarding response accuracy. Each binary sequence contained two events represented by two symbolic stimuli, specifically a square and a circle. The stimuli corresponded to response keys (“F” and “J,” respectively). Key assignment was counterbalanced among the participants. At the beginning of each session, a fixation cross appeared for 300 ms in the center of the display. In each trial, one of the stimuli, either a circle or a square, (2° × 2°) of visual angle appeared on the center of the display for 200 ms. When the stimuli were presented, the participants were instructed to press the corresponding key as quickly as possible without the cost of response accuracy. Participants were required to press a key within 1,000 ms. If no response was received, the trial was considered a no-response trial. Between each trial, participants were instructed to blink and maintain their focus on the blank screen for 2,000 ms (Figure 1A).

**Figure 1.**
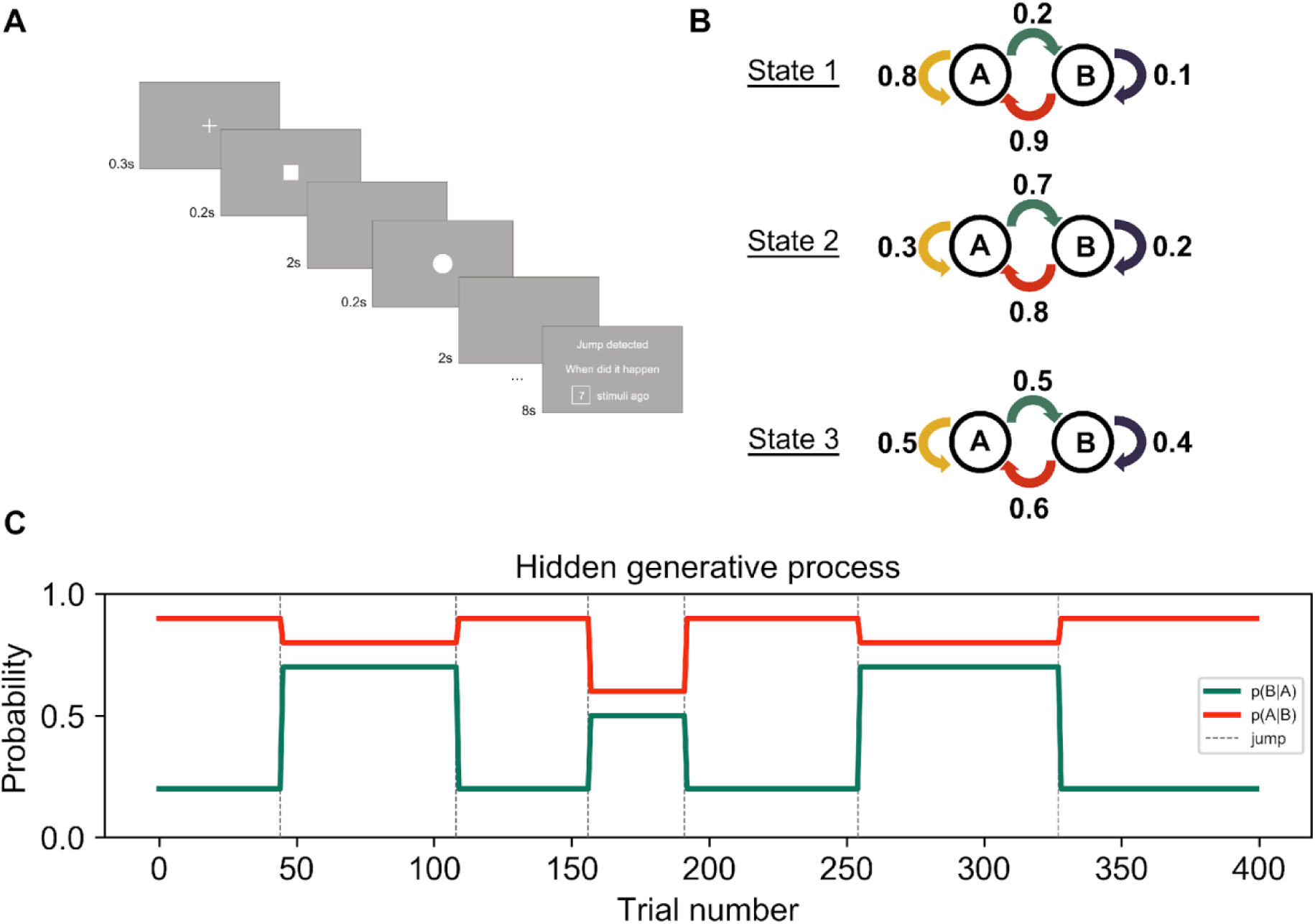
Experimental design. (A) Experimental design. (B) Event transition probabilities in each state. The first pair corresponds to the probability that Event A repeats, *p*(A|A), and the probability of transition to Event B, *p*(B|A), represented by the yellow and green arrows. The second pair corresponds to the probability of Event B repeating, *p*(B|B), and the probability of transition to Event A, *p*(A|B), represented by the purple and red arrows. (C) Evolution of hidden states across trials in one session. Lines indicate how *p*(B|A) and *p*(A|B) change across trials. The colors are the same as those in (B). Jumps are marked by dashed grey lines.

In the second task, participants were informed that the binary sequences contained predetermined event probabilities for each trial. When the participants observed that the event probabilities changed, they pressed the space bar, and a jump detection screen would appear. The event probabilities and jumps were not disclosed to participants and had to be inferred on the basis of the sequences. Participants were only informed that constant event probabilities may last for a period of time, in which was in the range of 30 to 70 trials. Participants had 8 s to report how many trials ago a jump had occurred. Their responses were displayed on the screen. Then, the participants continued with the TCRT task. Two practice sessions with 144 trials were conducted. During each practice session, verbal feedback was provided to familiarize the participants with jumps; no feedback was provided during the formal experiment. All the experimental materials are available (https://osf.io/6qpng/).

### Task Design

The binary sequence was generated from a Hidden Markov Model (HMM). HMM is one kind of probabilistic model that is able to generate a sequence with two stochastic processes where one involves the transition among different states while the other involves the emission of observations from a given state. HMM elements include the number of states in the model, the number of observations per state, the state transition probability, and the emission probabilities that determine what type of observation is generated in a particular state (Franzese et al., 2019; Rabiner, 1989).

Our HMM had three states: States 1–3. Two observations could be generated per state. The state transition probability, which determines how long one state would last, was 1/75. State transition probability can also be understood as the probability of a change between states, that is, a jump. The emission of observations was conducted by defining two pairs of *event transition probabilities*. Because binary sequences comprised two events, emission probabilities were grouped into two pairs, each with a probability of event alteration and a probability of event repetition. For two events A and B, the two pairs of event transition probabilities can be represented by *p*(*B*|*A*) and *p*(*A*|*A*) and *p*(*A*|*B*) and *p*(*B*|*B*). In State 1, *p*(*B*|*A*) was 0.2, and *p*(*A*|*A*) was 1 − 0.2 = 0.8; *p*(*A*|*B*) was 0.9, and *p*(*B*|*B* was 1 − 0.9 = 0.1. In State 2, *p*(*B*|*A*) was 0.7, and *p*(*A*|*A*) was 0.3; *p*(*A*|*B*) was 0.8, and *p*(*B*|*B*) was 0.2. In State 3, *p*(*B*|*A*) was 0.5, and *p*(*A*|*A*) was 0.5; *p*(*A*|*B*) was 0.6, and *p*(*B*|*B*) was 0.4 (Figure 1B). Different event transition probabilities result in different sequence statistics. The generated sequence in State 1 was dominated by one type of event, either squares or circles. In State 2, the generated sequence was more likely to have interleaving events. Because all event transition probabilities were close to chance in State 3, the generated sequence was random. Jumps indicated the changing of states and therefore the changing of the probabilities of event transition. An additional constraint was imposed to make jumps more noticeable; the change in the odds ratio for one of the event transition probabilities compared with the previous probability was at least four. As a result, only changes from State 2 to State 1, from State 3 to State 1, or from State 1 to State 1, State 2, or State 3 occurred. Each session consisted of a sequence with 400 trials generated using the model (Figure 1C).

State transition probability determined how probable jumps, that is, a change of state, were. A change of state meant a change in emission probability, with two pairs of event transition probabilities in each state. Different emission probabilities resulted in different sequence statistics. The fact that each state had two pairs of event transition probabilities was not disclosed to participants; they were only informed that they should keep track of the event probabilities.

### Bayes-optimal learning model

We used the Bayes-optimal learning model (<u>Meyniel et al., 2016</u>; <u>Meyniel, 2020</u>) (https://github.com/florentmeyniel/MinimalTransitionProbsModel). In the model, the Bayes theorem is used to infer posterior distributions for a given trial, represented by θ*_t_*, given a set of model assumptions *M* and previous observations *y*_1:*t*_.

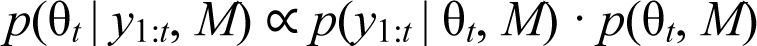

On the basis of information learned from the binary sequence, the posterior distributions at trial θ*_t_*differed.

1. θ*_t_* as event probability: If the participants learned the probabilities of the two events, then θ*_t_*would be one dimensional, *p*(*A*) and *p*(*B*). We referred to this model as *M*_EP_, a model of event probability.
2. θ*_t_* as event transition probability: If the participants learned the event transition probabilities that generated the observations, then θ*_t_* would be two dimensional, where one dimension corresponded to *p*(*B|A*) and *p*(*A|A*) and the other corresponded to *p*(*A|B*) and *p*(*B|B*). We referred to this model as *M*_ETP_, a model of event transition probability.

The Bayes-optimal learning model is further detailed in Section 1 of Supplementary Materials.

### Quantification of Surprise

Shannon Surprise (*I_S_*) was used to quantify surprise in each observation. It was calculated by taking the negative logarithm base 2 of the probability of the observation. Binary sequences with *n* trials consisted of two events, *E_t_* ∈ {*A*, *B*} for *t* = 1, 2, …, *n*. Surprise differed depending on whether θ*_t_*was event probability or event transition probability.

1. θ*_t_* as event probability:

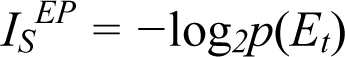

where *p*(*E_t_*) ∈ {*p*(*A*), *p*(*B*)} and *EP* stands for event probability. Surprise was the negative log likelihood of an event in a given trial. Studies (<u>Kolossa et al., 2013</u>, <u>2015</u>; <u>Higashi et al., 2017</u>) have defined surprise in relation to an event as predictive surprise, denoted by *I_S_^EP^*. Trial-by-trial predictive surprise values were computed on the basis of the trial-by-trial posterior distribution of *M*_EP_ (Figure 2A).

1. θ*_t_* as event transition probability:

**Figure 2.**
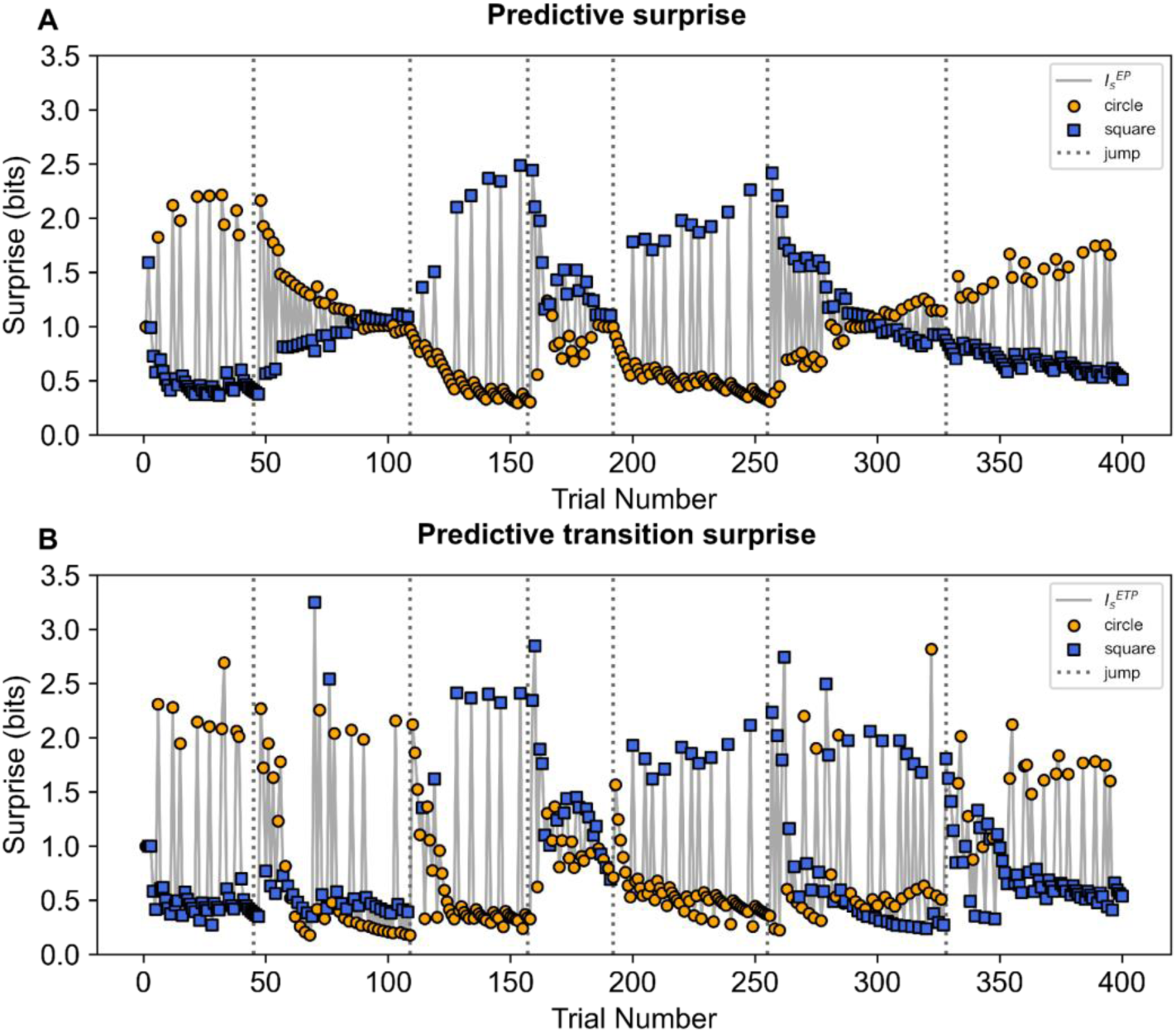
Trial-by-trial surprise values. (A) Predictive surprise values across trials. Predictive surprise denoted by I_S_^EP^ and represented by grey lines. The circle stimuli are represented by the orange circles, and the square stimuli are represented by the blue squares. Jumps are represented by dashed gray lines. (B) Predictive transition surprise values across trials. Predictive transition surprise is denoted by I_S_^ETP^ and plotted using grey lines.

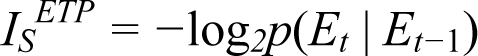

where *p*(*E_t_* | *E_t-1_*) ∈ {*p*(*B*|*A*), *p*(*A*|*A*), *p*(*A*|*B*), *p*(*B*|*B*)}, and *ETP* stands for event transition probability. Surprise was the negative log likelihood of an event transition in a given trial. In accordance with the method of <u>Higashi et al. (2017</u>), surprise calculated by accounting for event transitions was called “predictive transition surprise” (*I_S_^ETP^*). Trial-by-trial predictive transition surprise values were computed on the basis of the trial-by-trial posterior distribution of *M*_ETP_ (Figure 2B).

### Behavioral Statistical Analysis

One participant was excluded from the behavior analysis because of a response accuracy below 85% and a jump detection score below zero (Supplementary Materials Sections 2 and 3); Thirty-seven participants were included in the behavioral statistical analysis.

#### State and Event Transition Probability RTs

To identify a difference in RTs between the three states and their corresponding event transitions, we performed a two-way (3 × 4) repeated-measures analysis of variance (ANOVA) with two within-subjects factors, namely state (State 1, State 2, State 3) and four event transitions (*p*(*A*|*A*), *p*(*B*|*A*), *p*(*B*|*B*), *p*(*A*|*B*)) (Supplementary Materials Section 4). The analysis was performed after aligned rank transformation (ART) of the data (<u>Wobbrock et al., 2011</u>). This nonparametric method was applied because our data did not meet the assumptions of a factorial ANOVA (<u>Leys and Schumann, 2010</u>). The ART was performed in the R studio (Version 1.2.5033) with the *ARTool* package (<u>Kay and Wobbrock, 2016</u>). Simple main effect analysis was conducted when significant interactions were observed by modifying conditions on state type to examine the effects of transitions within each state. Post hoc tests were performed to identify differences between event transitions. In the post hoc tests, *p*-values were adjusted through Bonferroni correction to control alpha inflation due to multiple comparisons.

#### High-Versus Low-Surprise RTs

An analysis was performed to determine whether predictive surprise and predictive transition surprise affected RTs. Trials were divided into high- and low-surprise trials. Trials with surprise values equal to or greater than 1 were considered high surprise. Those with surprise values lower than 1 were considered low surprise. The value 1 was the dividing point because this value represents the point at which the observed likelihood is 50% (*−*log_2_(1/2) = 1).

A two-way (3 × 2) repeated-measures ANOVA with two within-subjects factors, namely state (State1, State 2, State 3) and surprise level (high, low), was performed after ART of the data because the assumptions of a factorial ANOVA were not met. Simple main effect analysis was performed on state to examine the effect of surprise in each state. The nonparametric Wilcoxon signed-rank test was used.

### Electroencephalogram Data

Electroencephalogram (EEG) data were recorded using 64 Ag/AgCl scalp electrodes mounted in an elastic cap in accordance with the international 10–20 system. The electrodes were Fp1/Z/2, AF7/3/4/8, F7/5/3/1/Z/2/4/6/8, FC5/3/1/2/4/6, FT7/FT8, C5/3/1/Z/2/4/6, CP5/3/1/Z/2/4/6, TP7/TP8, P7/5/3/1/Z/2/4/6/8, PO7/Z/8, and O1/Z/2. Six additional channels were used to measure eye movement and for reference in the offline analysis. To measure vertical eye movement, two electrodes were placed on the supraorbital and infraorbital ridges of the left eye; to measure horizontal eye movement, two electrodes were placed on the outer canthi of the left and right eyes. The other two electrodes (M1 and M2), serving as offline reference, were attached to mastoid sites. For all electrodes, impedance was maintained below 10 k . Online EEG data were recorded using the BrainAmp amplifier with a low-pass filter of 1,000 Hz. EEG data was recorded using a BrainVision Recorder (v0.23.0). The sampling rate was 1,000 Hz.

### EEG Data Preprocessing

EEG data were preprocessed using MNE software version 0.23.0 (<u>Gramfort et al., 2013</u>). Bad channels were identified through visual inspection. Continuous data were linearly detrended and bandpass filtered (0.1–30 Hz). Continuous data were then segmented into 1,200-ms epochs, with 200-ms prestimulus and 1,000-ms poststimulus intervals. We re-referenced each channel’s data with the average values of the M1 and M2 channels. Independent component analysis (ICA) was performed to identify signals related to blinking, eye movement, and muscle movement. These task-irrelevant signals were removed after ICA. Epochs were baseline corrected with the prestimulus interval. On the basis of each surprise array, predictive surprise, and predictive transition surprise, epochs were sorted into the following conditions: high surprise and low surprise, respectively for State 1, State 2, and State 3. Epochs with incorrect responses were excluded. Artifact rejection was performed when epoch signals in any channel exceeded ±75 μV. For bad channels, interpolation was performed to recover data. Grand average ERPs were obtained for epochs in all conditions.

#### Single-Trial P300 Estimates

We estimated single-trial P300 at the Pz electrode where the P300 was traditionally found to be maximal (<u>Duncan-Johnson and Donchin, 1977</u>). Single-trial P300 estimates were extracted on the basis of the grand average ERPs of all epochs after preprocessing. The time point of maximal modulation was identified, and single-trial data were extracted with a time window of ±60 ms (Barcelo et al., 2008; Mars et al., 2008; Kolossa et al., 2013). If this time point was one standard deviation away from the mean of the mean time for all participants, the participant’s data would be excluded from analysis. Two participants were excluded, and the sample size for the single-trial P300 analysis was 30 (mean = 384 ms, standard deviation [SD] = 28.8 ms).

### EEG Data Statistical Analysis

Six participants were excluded from the EEG data statistical analysis because of malfunction in the devices during data collection. Thirty-two participants were included in the final analysis.

#### Surprise: High-Versus Low-Surprise ERPs

For ERPs, the difference between high- and low-surprise conditions were examined. High- and low-surprise ERPs were compared, and an analysis was performed for each state. A nonparametric cluster-based permutation test (<u>Maris and Oostenveld, 2007</u>; <u>Jas et al., 2018</u>) was also performed. Clusters were considered significant if their *p*-values were below 0.05 (cluster-forming threshold *p* < 0.05, cluster-level *p* < 0.05, 5,000 permutations).

The cluster-based permutation test results were carefully organized to avoid misleading interpretations. The significance of effect latency and location was not fully controlled for in the statistics (<u>Sassenhagen and Draschkow, 2019</u>). For this reason, the results are reported in a descriptive manner.

### Single-Trial Analysis: Model Parameter Estimation and Model Evaluation

An analysis was performed to determine whether a model based on event transitions or a model based on event probabilities would better account for trial-by-trial RTs and P300. A hierarchical general linear model (GLM) was used for the analysis, and trial-by-trial predictive transition surprise *I_S_^ETP^*and predictive surprise *I_S_^EP^* were used as regressors. To estimate the parameters, each participant’s data were fitted with a slope and intercept, indicating how each individual’s trial-by-trial data changed as a function of surprise. To evaluate the models, variational free energy *F*, which is an approximation of the log-likelihood of data given a model log(*p*(y | m)) and not directly computable, was calculated (<u>Mars et al., 2008</u>; <u>Kolossa, 2016</u>). The Bayes factor (<u>Kass and Raftery, 1995</u>) was then computed to compare fit between the models, that is, the ratio between model likelihoods. If the natural logarithm is used, then the log-Bayes factor is the difference between the models’ likelihoods. Thus, for the two models, *M* = {*M*_ETP_, *M*_EP_}:

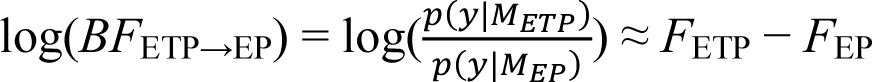

Positive values for log-Bayes factors would provide stronger evidence for the superiority of *M*_ETP_ over *M*_EP_, whereas negative values would provide stronger evidence for the superiority of *M*_EP_. The spm_PEB.m function was used to estimate the models’ parameters and evaluate the models. This function can be found in the Statistical Parametric Mapping 12 package in Matlab. The hierarchical GLM was used in accordance with methods used in other studies (<u>Mars et al., 2008</u>; <u>Kolossa et al., 2013</u>; <u>Kolossa, 2016</u>). Here, only a brief summary of this model is provided. A more detailed explanation is provided in Supplementary Materials Section 5.

## Results

### Behavioral Results

#### State and Event Transition Probability RTs

The main effect of state (*F*_2, 72_ = 4.62, *p* = 0.01, *η*^2^ = 0.11) and the main effect of event transition (*F*_3, 108_ = 16.45, *p* < .001, *η*^2^ = 0.31) were both significant. The interaction between state and event transition (*F*_6, 216_ = 53.26, *p* < .001, *η*^2^ = 0.60) was also significant. Simple main effect analysis showed significant differences in event transitions in State 1 (χ^2^ = 30.63, *p* < .001, degrees of freedom [df] = 3) and State 2 (χ^2^ = 11.06, *p* = 0.01, df = 3) (Figure 3A). Post hoc tests showed significant differences in RTs between both pairs of event transitions in State 1 For the first pair, the RT of *p*(*A*|*A*) = 0.8 was shorter than that of *p*(*B*|*A*) = 0.2 (*median_p(A|A)_=0.8* = 394 *ms*, *median_p(B|A)=0.2_* = 498 *ms*, *p* < .001, *r* = 1.06). For the second pair, the RT of *p*(*B*|*B*) = 0.1 was longer than that of *p*(*A*|*B*) = 0.9 (*median_p(B|B)=0.1_* 469 *ms*, *median_p(A|B)_*=0.9 = 424 *ms*, *p* < .001, *r* = −0.64). The RT of *p*(*A*|*A*) was also shorter than those of *p*(*B*|*B*) (*median_p(A|A)=0.8_* 394 ms, *median_p(B|B)=0.1_* = 469 *ms*, *p* < .001, *r* = 0.91) and *p*(*A*|*B*) (*median_p(A|A)=0.1_* = 394 *ms*, *median_p(A|B)_=0.9* = 424 *ms*, *p* < .001, *r*= 0.76). The RT of *p*(*B*|*A*) was also longer than those of *p*(*B*|*B*) (*median_p(A|A)=0.8_ p* < .001, *r* = −0.58) and *p*(*A*|*B*) (*median_p(A|A)=0.8_ p* < .001, *r* = −1.05). Post hoc tests showed significant differences in RTs between both pairs of event transitions in State 2. For the first pair, the RT of *p*(*A*|*A*) = 0.3 was longer than that of *p*(*B*|*A*) = 0.7 (*median_p(A|A)=0.3_* = 475 *ms*, *median_p(B|A)_*=0.7 = 430 *ms*, *p* < .001, *r* = −0.92). For the second pair, the RT of *p*(*B*|*B*) = 0.2 was longer than that of *p*(*A*|*B*) = 0.8 (*median_p(A|A)=0.2_* = 454 *ms*, *median_p(A|B)=0.8_* = 428 *ms*, *p* < .001, *r* = −0.50). The RTs of *p*(*A*|*A*) were also longer than those of *p*(*B*|*B*) (*median_p(B|B)=0.3_* = 475 *ms*, *median_p(B|B)=0.8_* = 454 *ms*, *p* = 0.02, *r* = −0.38) and *p*(*A*|*B*) (*median_p(A|A)=0.3_* = 475 *ms*, *median_p(A|B)=0.8_* = 428 *ms*, *p* < .001, *r* = −0.92). The RT of *p*(*B*|*A*) was shorter than that of *p*(*B*|*B*) (*median_p(B|A)=0.7_* = = 430 *ms*, *median_p(B|B)=0.2_* = 454 *ms*, *p* = .001, *r* = 0.52).

**Figure 3.**
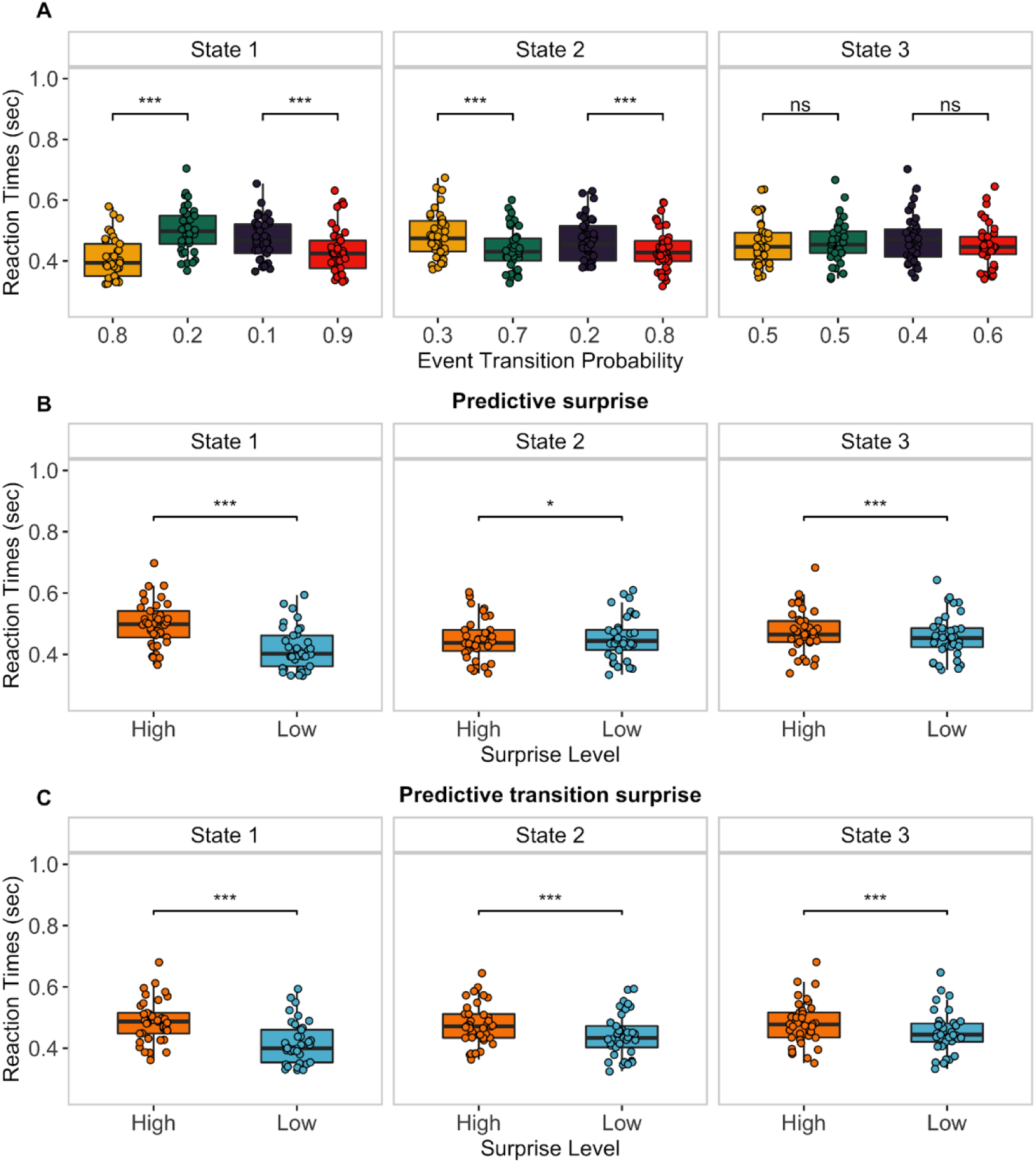
Behavior results. (A) RTs in each state by surprise level. Event transition probabilities in each state a on the X-axis. Only the RTs for each pair of event transitions are plotted. The rest of the results can be found in the main text. The colors corresponded to the colors of the arrows in Figure 1B. (B) RTs between high and low predictive surprise in each state. (C) RTs between high and low predictive transition surprise in each state. (***: *p* < .001, **: *p* < .01, *: *p* < .05, ns: nonsignificant).

#### Predictive Surprise: High-Versus Low-Surprise RTs

The main effect of state (*F*_2, 72_ = 11.98, *p* < .001, *η*^2^ = 0.25) and the main effect of surprise type (*F*_1, 36_ = 67.59 *p* < .001, *η*^2^ = 0.65) were significant. The interaction between state and surprise level (*F*_2, 72_ = 91.17, *p* < .001, *η*^2^ = 0.72) was also significant. Simple main effect analysis showed the effects of surprise in all three states (State 1: high surprise *median* = 498 ms, low surprise *median* = 402 ms, *p* < .001, *r* = −1.11; State 2: high surprise *median* = 437 ms, low surprise *median* = 444 ms, *p* = 0.03, *r* = 0.36; State 3: high surprise *median* = 465 ms, low surprise *median* = 453 ms, *p* < .001, *r* = −0.55; Figure 3B).

#### Predictive Transition Surprise: High-Versus Low-Surprise RTs

The main effect of state (*F*_2, 72_ = 11.63, *p* < .001, *η*^2^ = 0.24) and the main effect of surprise level (*F*_1, 36_ = 90.24, *p* < .001, *η*^2^ = 0.71) were significant. The interaction between state and surprise level (*F*_2, 72_ = 35.03, *p* < .001, *η*^2^ = 0.49) was also significant. Simple main effect analysis showed the effects of surprise in all three states (State 1: high surprise *median* = 487 ms, low surprise *median* = 400 ms, *p* < .001, *r* = −1.11; State 2: high surprise *median* = 471 ms, low surprise *median* = 433 ms, *p* < .001, *r* = −0.92; State 3: high surprise *median* = 478 ms, low surprise *median* = 444 ms, *p* < .001, *r* = −0.82; Figure 3C).

### EEG Results

#### Predictive Surprise: High-Versus Low-Surprise ERPs

Cluster-based permutation tests showed significant differences between high- and low-surprise ERPs in States 1 and 2 (Figure 4A). Clusters exhibiting characteristics of the P300 were observed. In State 1, the temporal profile of the cluster was from 329 to 642 ms (cluster *p* value < .001). In State 2, the temporal profile of the cluster was from 249 to 563 ms (cluster *p* value = 0.02). In State 3, no difference between the high- and low-surprise ERPs was observed.

**Figure 4.**
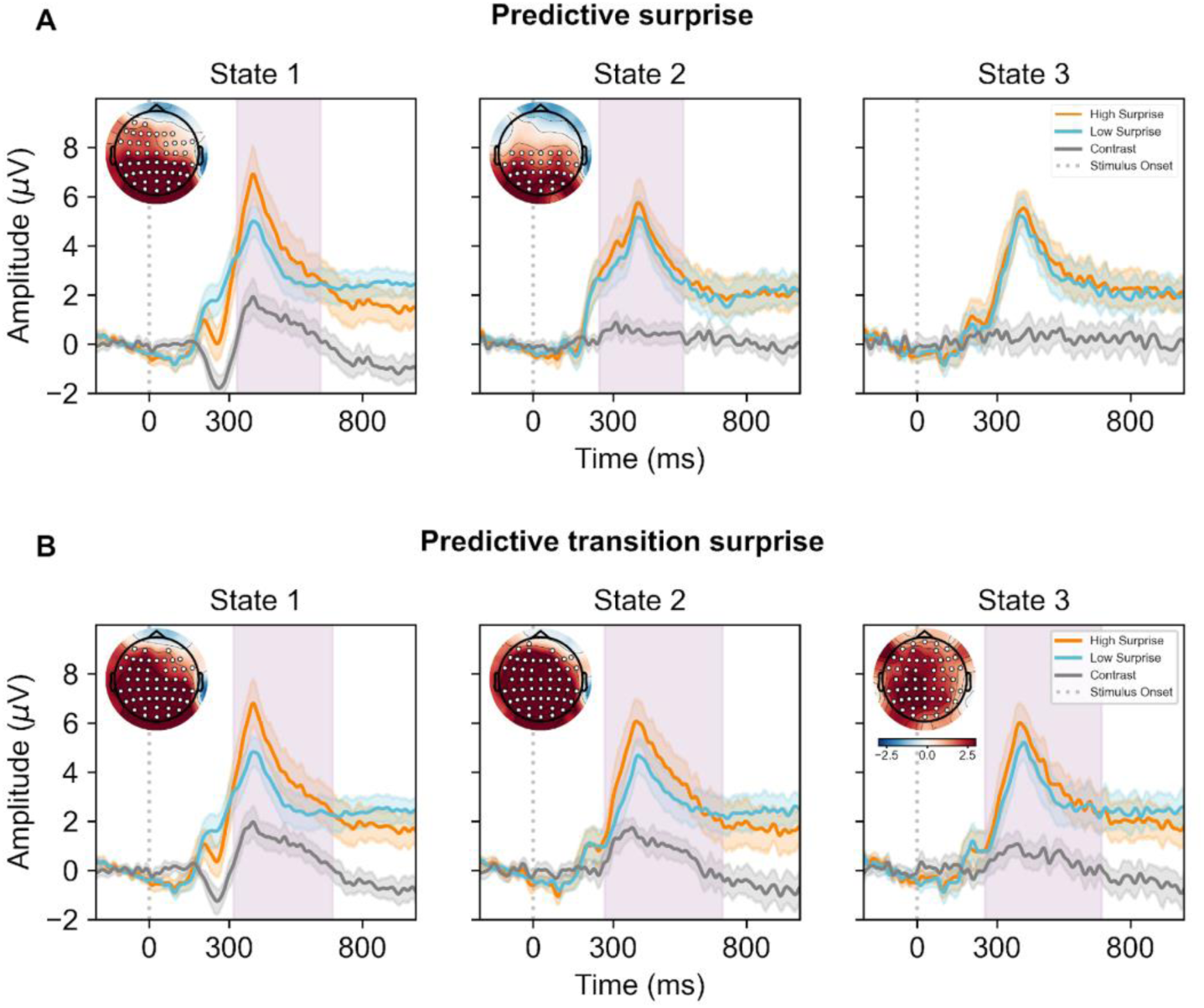
EEG ERPs results. (A) Difference in ERPs between high and low predictive surprise in each state. High-surprise ERPs across all participants are represented by orange lines. Low-surprise ERPs across all participants are represented by blue lines. The contrast between high and low predictive surprise is represented by grey lines. Shaded areas around the lines indicate confidence intervals. The light thistle shaded area represents the temporal profile of the clusters. The topographical map on the upper-left part of each panel represents averaged *t* values for a cluster within 300–450 ms. Electrodes with significant differences are marked by white circles. (B) Difference in ERPs between high and low predictive transition surprise in each state.

#### Predictive Transition Surprise: High-Versus Low-Surprise ERPs

Cluster-based permutation tests showed significant differences between high- and low-surprise ERPs in all three states (Figure 4B). Clusters exhibiting characteristics of the P300 were observed in all three states. In State 1, the temporal profile of the cluster was from 318 to 688 ms (cluster *p* value < .001). In State 2, the temporal profile of the cluster was from 268 to 709 ms (cluster *p* value < .001), and in State 3, the temporal profile of the cluster was from 255 to 692 ms (cluster *p* value < .001).

### Results of Single-Trial Model-Based Analysis

Table 1 presents the single-trial RTs and the P300 results from the model-based analysis 1. These results were obtained by using the hierarchical GLM described in the “Methods” section. For both RTs and the P300, the model fitting accuracy favored *M*_ETP_ over *M*_EP_, suggesting that trial-by-trial variation in RTs and the P300 are better explained by predictive transition surprise than by predictive surprise. These results indicate that increased RTs and the P300 are observed when more surprising events occur. All the processed data is available (https://osf.io/6qpng/).

**Table 1.** Log-Bayes factors between *M*_ETP_ and *M*_EP_

| Table 1. Log-Bayes factors between $M_{ETP}$ and $M_{EP}$ | |
| --- | --- |
| $\ln(BF_{M_{ETP} - M_{EP}})$ | |
| RTs | 266 |
| P300 | 36 |

## Discussion

We examined the P300 in relation to sequence knowledge in changing environments. Participants were presented with a changing and probabilistic sequence generated by an HMM. A model of event probability and a model of event transition probability were examined to determine which better captured participants’ behavior. In each state, RTs changed depending on event transition probability. More probable event transitions were reacted to faster, whereas less probable event transitions were reacted to slower. The model-based analysis revealed that predictive transition surprise better explained RTs and the P300 overall. High-surprise trials were associated with longer RTs and larger P300 amplitudes in all three states. Regression analysis indicated that predictive transition surprise better predicted trial-by-trial RTs and the P300 than did predictive surprise. We observed that the state of the environment influenced the representation of sequence knowledge in the brain. In addition, the P300 may have reflected the sequence knowledge of the probability of transition between events in the changing environment.

This study explored how sequence knowledge changes in a dynamic environment, as have other studies (<u>Mars et al., 2008</u>; <u>Kolossa et al., 2013</u>; <u>Higashi et al., 2017</u>). Larger P300 amplitudes were observed in high predictive transition surprise conditions across all three states. Larger P300 amplitudes were also observed in high predictive surprise conditions in States 1 and 2 but not in State 3, possibly because States 1 and 2 have more number of trials before a change occurs; this may have created a stable state environment similar to a block design, in which sequence knowledge of event probability may be useful. As a result, predictive surprise exhibited strong performance in States 1 and 2. This study did not fully control the frequency of the changes in the hidden states of States 1 and 2, and further research is required to determine the extent to which this would affect encoded sequence knowledge.

Our results support the hypothesis that event transition probability serves as a building block of sequence knowledge (<u>Meyniel et al., 2016</u>). <u>Meyniel et al. (2016)</u> proposed the event transition model that accounted for simulated behavioral results from other studies (<u>Huettel et al., 2002</u>; <u>Cho et al., 2002</u>; <u>Squires et al., 1976</u>) with different sequence designs and across different experimental modalities including behavioral, functional magnetic resonance imaging, and EEG studies. Another study from the same group showed that the inference of event transition probability was independent of the types of sequence statistics. The event transition model performed better for all sequence statistics, including fully stochastic, frequency-biased, alternation-biased, and repetition-biased sequences (<u>Maheu et al., 2019</u>).

Our study examined whether the state of the environment affects encoded sequence knowledge, and the results indicate that the event transition model performed better for dynamic sequences concatenated with different types of sequence statistics. Thus, our results alongside those of other studies (<u>Meyniel et al., 2016</u>; <u>Maheu et al., 2019</u>) demonstrate that representation is based on event transition probability, regardless of the type of sequence statistics and whether the sequence statistics change over time.

Our participants were only asked to keep track of event probabilities. Participants identified hidden regularities without being instructed to do so. Their RTs indicated dependence on event transition probabilities. This result is related to key concepts in statistical learning about the ability to identify structures and extract regularities in the environment (<u>Schapiro and Turk-Browne, 2015</u>; <u>Conway, 2020</u>). Humans have found to possess this ability in various domains and modalities, including language acquisition (<u>Saffran et al., 1996</u>) and the processing of nonlinguistic auditory tones (<u>Saffran et al., 1999</u>), visual objects (<u>Fiser and Aslin, 2002</u>), and visual scenes (<u>Brady and Oliva, 2008</u>). It is thus suggested as a domain-general ability, and similar computational mechanisms may operate across multiple neural networks (<u>Frost et al., 2015</u>). The participants’ ability to identify hidden regularities in streams of events and input information highlights the brain’s need to create accurate representations of the environment, especially for changing environments, for which knowledge must be constantly updated. Our results suggest that humans can identify temporal statistics regardless of whether they are explicitly given.

This study has several limitations. The first is the design of the TCRT task, which could not determine whether the observed P300 was caused by surprise due to visual stimuli or surprise related to the motor responses, as other studies have described (<u>Mars et al., 2008</u>; <u>Kolossa et al., 2013</u>; <u>Higashi et al., 2017</u>). The second limitation is that the scope was limited to binary sequences. Whether the results can be generalized to complex sequences with more events remains unclear. Finally, <u>Higashi et al. (2017)</u> performed a single-trial analysis that revealed that predicted surprise better explained P3b and that predictive transition surprise better explained P3a. The FCz electrode was selected for P3a, and the Pz electrode was selected for P3b because these electrodes were most reported to represent the corresponding P300 sub-components (<u>Kolossa et al., 2015</u>). The relationship between sequence knowledge and these sub-components of the P300 in a changing environment should be investigated. However, we did not fully control the frequency of changes in the hidden states, and for this reason, our experimental design may not be ideal for such an investigation. This factor should be considered in future investigations of whether each sub-component of the P300 reflects a different type of sequence knowledge in a changing environment.

We performed the model-based analysis to study the P300 in a changing environment. Our results indicate that fluctuations were better explained by predictive transition surprise, suggesting that humans learn the event transition probabilities and that the P300 may be linked to this type of sequence knowledge. The inference of event transitions provided additional resolution, that is, temporal event contingencies, which not only facilitate better representation of the environment but also enable the accurate prediction of events.

## Acknowledgements

This work was supported by grants from National Science and Technology Council (110-2410-H-038-009, 111-2410-H-038-009-MY2) to T.Y.H.

## Author Contributions

K.P.C: Conceptualization, Methodology, Software, Formal analysis, Investigation, Writing – Original Draft, Visualization. T.Y.H.: Conceptualization, Methodology, Resources, Writing – Review & Editing, Visualization, Supervision, Project administration, Funding acquisition

## Supplementary Materials

### 1. Bayes-optimal learning model

In the Bayes-optimal learning model (Meyniel, et al., 2016; Meyniel, 2020), Bayes’ theorem is used to infer the posterior distributions of any given trial, denoted by θ*_t_*, given a set of model assumptions *M* and previous observations *y*_1:*t*_.

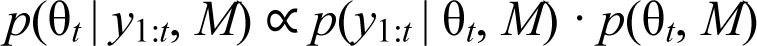

The model assumptions *M* are related to how we implemented the HMM. First, state transition probability *p*_c_ was given, and because the moments in which the state would change were not disclosed to the participants, their possibility in any given trial should be considered. Second, the model assumed that when a change occurred, the posterior distributions would be resampled in the range of 0 to 1. The model was not aware of the constraint mentioned in the main text, which ensured that the change in odds ratio for one of the event transition probabilities compared to the previous would be at least four. Finally, a key aspect of the HMM is the Markov property; if θ*_t_*_−1_ is known, then the previous observation *y*_1:*t*−1_ is not needed to estimate θ*_t_*; thus, the term *p*(θ*_t_* | *y*_1:*t*_, *M*) is rewritten as follows:

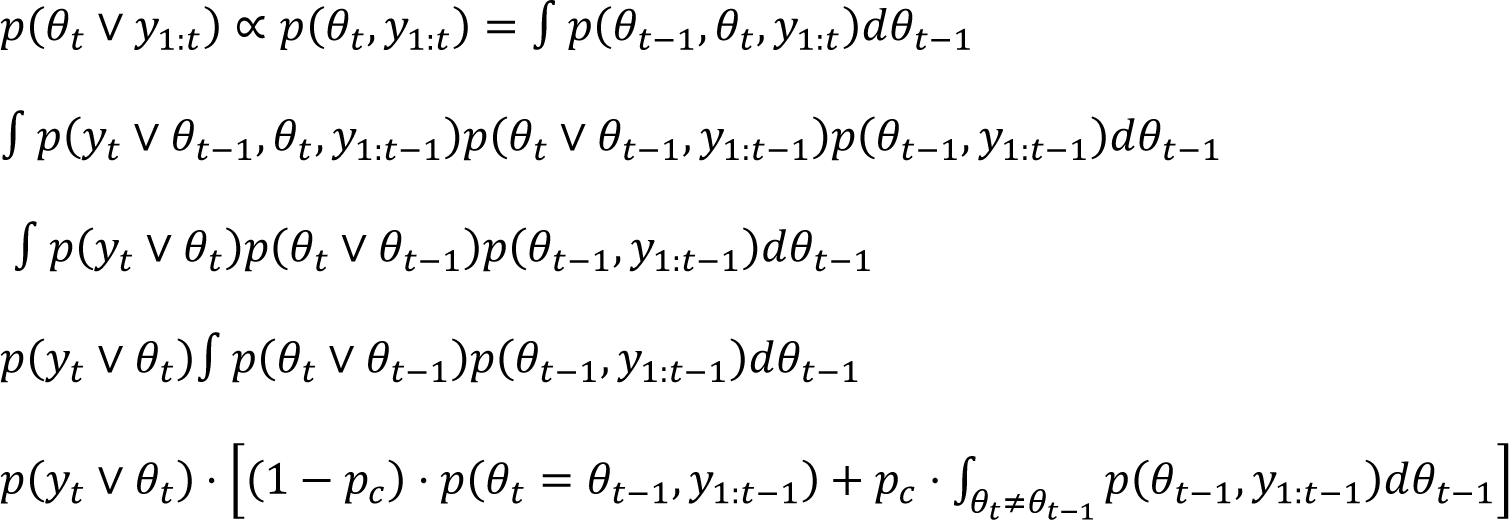

Joint distribution, *p*(θ*_t_, y*_1:*t*_) was calculated recursively by starting from a prior observation and updating the distribution at each time point. The prior about θ*_t_* was flat. The integral was computed by using a probability grid for numerical integration across trials. Finally, we obtained the posterior distribution by normalizing the joint distribution.

(1) θ*_t_* as event probability: If participants learned the probabilities of the two events, then θ*_t_* would be one dimensional, *p*(*A*) and *p*(*B*). The probability grid in this case would consist of 100 values.

(2) θ*_t_* as event transition probability: If the participants learned the event transition probabilities that generated the observations, then θ*_t_* would be two dimensional, where one dimension corresponded to *p*(*B|A*), *p*(*A|A*) and the other corresponded to *p*(*A|B*), *p*(*B|B*). The probability grid in this case would be two dimensional, with 100 × 100 values, one dimension corresponding to *p*(*B|A*), *p*(*A|A*), and the other corresponding to *p*(*A|B*), *p*(*B|B*).

Trial-by-trial posterior probability distributions were calculated using the codes provided by Meyniel, et al. (2016) online. (https://github.com/florentmeyniel/MinimalTransitionProbsModel).

### 2. Jump Detection Task Analysis

This analysis was performed to quantify the participants’ performance in the jump detection task by using the informedness index, which is calculated by dividing the hit rate by the false alarm rate (Meyniel, et al., 2015). Values exceeding zero indicate detection above chance, and values below zero indicate detection below chance. If an estimated jump was within ±10 trials of an actual jump, it was considered a hit. If no estimated jump was within this range, it was considered a miss. The hit rate was calculated by dividing hit counts by the sum of hit and miss counts. If an estimated jump was outside this range, it was counted as a false alarm. The false alarm rate was calculated by dividing the false alarm count by the total number of states in the sequence.

For each session, a Wilcoxon signed-rank test was conducted on the informedness index to determine whether the median was significantly higher than zero. In addition, a Wilcoxon signed-rank test was conducted to identify differences between sessions.

### 3. Response Accuracy and Performance in Jump Detection Task

One participant was excluded from the behavior analysis because of a response accuracy below 85% and a jump detection score below zero. For the rest of the participants, the response accuracy was 97% (SD = 0.03). For the jump detection task, the median of the informedness index was significantly higher than zero for both Session 1 (*median* = 0.69, SD = 0.38, *p* < .001, *r* = 0.83) and Session 2 (*median* = 0.88, SD = 0.23, *p* < .001, *r* = 0.88). A significant difference between Sessions 1 and 2 was observed (*p* = .003, *r* = 0.48), suggesting that Session 2 had a learning effect. Overall, these analyses revealed that the participants were attentive and aware of jumps during the task.

### 4. RTs Preprocessing

Before we investigated differences in RTs between the three states and their corresponding event transition probabilities, the RTs data were preprocessed. RTs for each state were divided into four event transitions. Trials were excluded from statistical analysis if they had incorrect responses or any responses that were above or below 3 SDs of the mean RT for each state, which were considered outliers. The first trials after the jump detection task were also excluded. We combined mean RTs from all participants in each RT distribution for the group analysis.

### 5. Hierarchical GLM

Data from all *S* participants were concatenated in a three-level GLM. We used the same notation as Kolossa, et al. (2013) and structured the three-level GLM as follows:

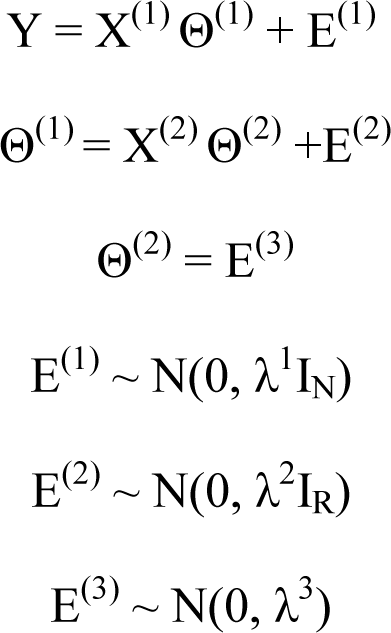

The first level of the GLM models single-trial data *Y* as a function of surprise values *X*^(1)^ and error *E*^(1)^. *Y* is a column vector containing data from all *S* participants. Thus, *Y* = [*y_s_*_=1_, *y_s_*_=2_, …*y_s_*_=*S*_]*^T^*, *Y* ∈ ℝ*^N^*, where *N* = ∑ *^S^ N*, which is the total number of trials from all participants, and []*^T^* indicates transposition. This model can handle unbalanced input data. For each subject, only RT data from trials with correct responses were included. For P300 responses, only trials with correct responses that were not rejected during EEG preprocessing were included in the analysis. *X*^(1)^ is the first-level design matrix that is block diagonal, with dimensions of *N* × *RS*, where *R* is the number of regressors for each subject. The regressors were explanatory variables, and we used *R* = 2 regressors; the first column was all ones, and the second column comprised model-dependent surprise values *I_S_^ETP^* or *I_S_^EP^*. To estimate parameters in the first level, Θ^(1)^, a matrix containing participant-specific ϑ ^(1)^ and θ_s_ ^(1)^, was used, with Θ^(1)^ = [ϑ*_s_*_=1_^(1)^, θ*_s_*_=1_^(1)^, ϑ*_s_*_=2_^(1)^, θ*_s_*_=2_^(1)^, …, ϑ*_s_*_=*S*_^(1)^, θ*_s_*_=*S*_^(1)^]*^T^*. This can be understood as the participant-specific intercept and slope to the surprise regressor.

The second level models the between-subjects differences, where first-level participant-specific parameters Θ^(1)^ are modeled as the variation in group parameters Θ^(2)^∈ ℝ^2^. The second-level design matrix *X*^(2)^ was formed by stacking up identity matrix *Ι*_2_∈ ℝ^2⨉2^ *S* times. To estimate parameters in the second level, Θ^(2)^ was used to represent group parameters Θ^(2)^ _= [ϑ_ (2)_, θ_ (2)_]_*T*.

The third-level design matrix *X*^(3)^ is an all-zero design matrix. This sets an unconstrained prior (Friston, et al., 2007; Kolossa, 2016) on the second-level parameters, thus making Θ^(2)^ = *_e_*(3).

Unknown parameters Θ in each level and model were estimated using parametric empirical Bayes (PEB) schemes (Friston, et al., 2002). The hierarchical model was collapsed into a single level and then augmented to estimate the unknown parameters and error in all levels by using the expectation maximization (EM) algorithm (<u>Kolossa, 2016</u>). Error in all levels was assumed to be normally distributed, with zero means and isotropic (i.e., spherical) covariances (<u>Kolossa et al., 2013</u>; <u>Kolossa, 2016</u>). Thus, identity matrices *I_N_* and *I_R_* were created, and trials (Level 1) and parameters (Level 2) were assumed to be independent from each other. To control the noise at each level, covariances in each level were parameterized as hyperparameters λ^1^, λ^2^, and λ^3^ (<u>Mars et al., 2008</u>). They are called “hyperparameters” because their values also affect parameters Θ to be estimated. The PEB schemes involve the use of EM to compute the posterior densities of the unknown parameters and the hyperparameters in each level. The posterior densities are point estimates and can be interpreted as given data representing the reliability of a parameter (<u>Mars et al., 2008</u>); in our case, this would indicate the effect of surprise on single-trial RTs and P300.

